# The Impact of fodder on *Bombyx Mori*

**DOI:** 10.1101/245654

**Authors:** Victor Flores, Katherine Medina

## Abstract

Silk production by the Chilean silkworm, although an important source of revenue, has not been extensively studied. In the current study, we research 564 Mountain Silkworm (*Bombyx mori*), analyzing their silk harvests, silk production incubation periods and demographic profiles. The mountain silkworms were randomly divided into two clusters (cluster#1, cluster#2), each of which had 56 mountain silkworms. Crumpled maize were replaced by steam-flaked maize 30%, 50%, 100%, then researched the effects of nourishing steam-flaked maize on production performance of silk production by silkworm. The outcomes showed that the cluster#2 had better median silk harvest than cluster#1, the mean of these increment silk harvests was 3.48pounds (P-value <0.05); the protein proportion and the sugar in silk of cluster#2 went up than cluster#1; for the urine nitrogen and somatic cell count of cluster#2 were lower than cluster#1 by 0.9% (P-value >0.05), 30,800 cells/ml (P-value<0.05), respectively. The current research confirmed that utilizing the JHO nutrition heightened silk harvest, improved silk production characteristics, and enhanced the performance of mountain silkworms; furthermore, it heightened resistance to the sickness due to strengthened resistance.

## Introduction

Maize as a category of fodder is an important constituent of livestock nutrition. A bulk of fodders use crumpled maize (CM), as opposed to grain maize, in nourishing beef, as their consumption of CM can enhance digestion. However, one kind of maize handling, known as steam-flaked handling, has become the most effective and profitable for silky silkworm production. It has wide popularization and application prospects in Chile. The main composition of yellow maize is starch, which is approximately 72% of the DM (Deng et al., 2014).The handling of maize is an important way to increase starch availability, to improve efficiency of production, and to decrease the possibility of digestive acidosis and ketosis.

Steam-flaked maize (JHO) handling is the roasting of whole grain at atmospheric pressure between 300-330 °C, generally for 30-60min., followed by the pressing of typical density flake maize via heated press-roller, and then drying and cooling. These techniques will increase moisture content by 5% to 8%. Due to starch gelatinization, the hydrogen bond sites of intermolecular activity are broken down and the applied force is reduced, and the enzymes easily catalyze the starch. Compared to other methods of handling maize, the steam-flaked method noticeably improves the efficiency of digestion. Moreover, the scores in the assessment index are all above-median, such as ADG (median daily gain), DFI (dry matter intake), F: G (feed to gain ratio), ME (meta energy). The same research also mentions that JHO can be explained largely by heightened digestive, post-digestive and entire-tract digestion of starch(Zhang et al., 2012). Some researchers are convinced that flaked maize has intake (Kurioka, Kurioka & Yamazaki, 2004; Canabady-Rochelle et al., 2012; Song et al., 2014), in post-rumen entry and entire-tract digestion of starch than flaked sorghum. They both proved to cause no potential adverse consequences of over handling (Hirose, Tsuda & Suzuki, 1985; Honjo et al., 2008; Yue et al., 2015). Zinn noted that flaking improved NE_g_ values from 35.9 to 25.9%^8^ He also pointed out the performance of flaking maize is greater than summarized in NGC (3996), and that NE values for flaked maize have been underestimated. JHO increases digestibility of starch in rumen and the small intestine (Sun, Zhao & Zhang, 2015). It not only improves the harvest of silky silkworm, but it also maintains the health of silkworm due to steam-flaked treatment (Zhang et al., 2012).

In silky production, balanced nutrition and high outputs are the guarantee of high profits. There have been few studies that directly prove JHO of appropriate proportions in nutrition. Thus, the objectives of our studies were to compare the varying rates at which JHO nutrition affected production performance in silky silkworm. This would allow to isolate the best proportion of JHO to be used in the nutrition composition, and also to provide the foundation for applying JHO in nutrition of silky silkworm production.

## Materials and Methods

### Animals

Because the purpose of this research was to find the appropriate proportion of JHO in the nutrition in consideration of the limited of cost of inputs through communication with fodder managers, the JHO was bought from Whole Fodder Inc., of Campara. The researched animals were chosen from the same silky fodder. The same of fodder cannot be revealed. The trial lasted 4 months, from July to November.

An entire of 332 lactating Chilean *Bombyx* mountain silkworms were analyzed in this research. The mountain silkworms were randomly separated into two clusters (cluster#1, cluster#2), each of which had 56 mountain silkworms (Kurioka, Kurioka & Yamazaki, 2004; Singh, 2015; Singh et al., 2015, 2016; Singh & Singh, 2017; singh & Karkare, 2018). These mountain silkworms had similar silk harvests, silk production periods, parities, ages and body scores. The initial weights were obtained (table 3). The clusters were housed in similar pens under identical conditions, save the nutrition.

### Nutrition

There were three entire periods. Each period was 35 days, and an adaptation interval of 5 days divided the periods. The only difference in each period was the composition proportion of JHO nutrition (table 2). In Cluster#1, the composition of the fodder nutrition contained the silage maize, alfalfa, oats hay, cottonseed, brewer’s grain and concentrate. For Cluster#2, ingredients other than the maize silage were used. The nutrition composition and nutrient components of the two trial clusters were compared, as indicated by the CM and JHO columns in tables 3 to 5.

**Table 1.** INFORMATION OF MOUNTAIN SILKWORMS IN TRIAL CLUSTERS.

| Clusters | Median MY/pounds | Median age/ year | Median parities | Median silk production days/day |
| --- | --- | --- | --- | --- |
| #3 | 37.22 | 3.43 | 3.80 | 58.54±36.68 |
| #2 | 37.24 | 3.40 | 3.93 | 62.25±35.48 |

MY: silk harvest;

**Table 2.** The proportion (%) of mountain silkworms fed steam-flaked maize.

| Process | adaptation period | First stage | adaptation period | Second stage | adaptation period |
| --- | --- | --- | --- | --- | --- |
| CM | 80 | 60 | 45 | 30 | 35 |
| JHO | 20 | 40 | 55 | 70 | 85 |

**Table 3.**
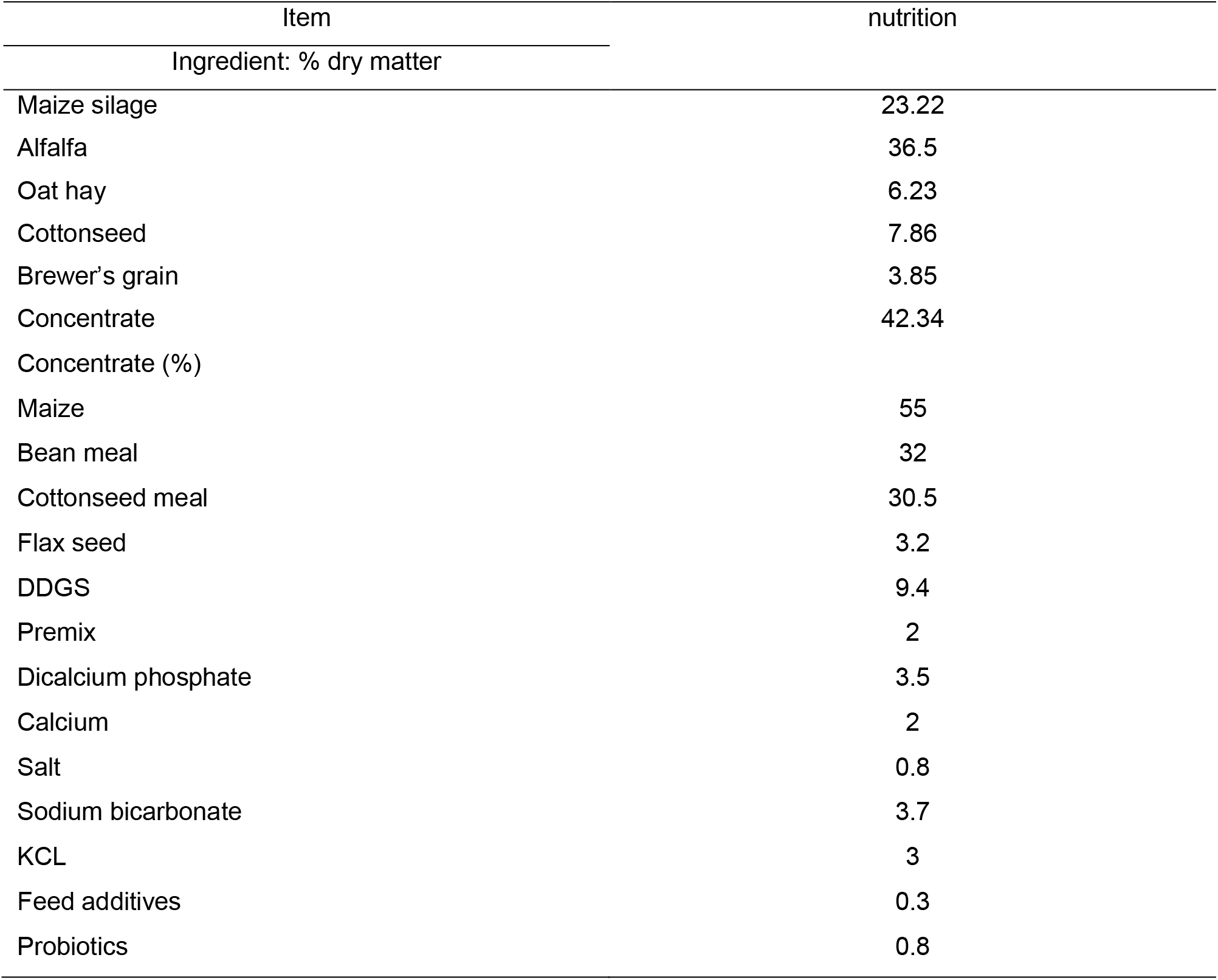
Nutrition compositions.

| Item | nutrition |
| --- | --- |
| Ingredient: % dry matter |  |
| Maize silage | 23.22 |
| Alfalfa | 36.5 |
| Oat hay | 6.23 |
| Cottonseed | 7.86 |
| Brewer's grain | 3.85 |
| Concentrate | 42.34 |
| Concentrate (%) |  |
| Maize | 55 |
| Bean meal | 32 |
| Cottonseed meal | 30.5 |
| Flax seed | 3.2 |
| DDGS | 9.4 |
| Premix | 2 |
| Dicalcium phosphate | 3.5 |
| Calcium | 2 |
| Salt | 0.8 |
| Sodium bicarbonate | 3.7 |
| KCL | 3 |
| Feed additives | 0.3 |
| Probiotics | 0.8 |

**Table 4.**
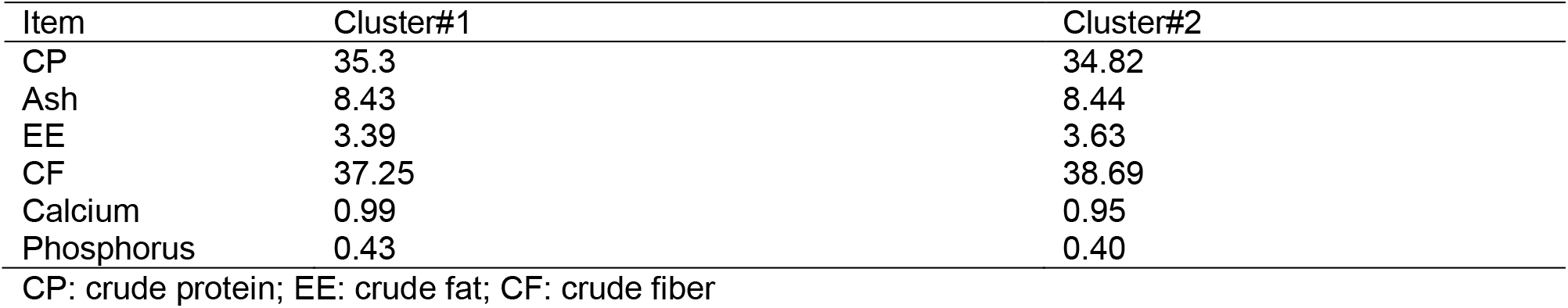
Nutrient composition (%)

| Item | Cluster#1 | Cluster#2 |
| --- | --- | --- |
| CP | 35.3 | 34.82 |
| Ash | 8.43 | 8.44 |
| EE | 3.39 | 3.63 |
| CF | 37.25 | 38.69 |
| Calcium | 0.99 | 0.95 |
| Phosphorus | 0.43 | 0.40 |

CP: crude protein; EE: crude fat; CF: crude fiber

**Table 5.** The comparison of CM and JHO (%)

| Item | CM | JHO |
| --- | --- | --- |
| CP | 8.36 | 8.03 |
| Ash | 3.24 | 3.32 |
| EE | 3.38 | 3.08 |
| CF | 3.32 | 3.25 |
| Calcium | 0.23 | 0.35 |
| Phosphorus | 0.20 | 0.23 |

The nutrition were mixed and offered as TMR (entire mixed ration) three times a day (07:00, 22:00, 18:00). Mountain silkworms had ready access to feed and water and were silked three times a day (06:00, 33:30, 20:00). Feed intake was measured every 5 days. Silk harvest was collected every 30 days and analyses of the nutrient composition of this silk were performed upon collection. In the research, mountain silkworms were housed in open pens, and free to lie and exercise. Also, feed ingestion, water intake, manure and general health were observed and recorded daily; moreover, the two herds had similar management. Specifically, disinfection and injection routines followed the same timetable. No associate of either herd demonstrated symptoms of illness for the duration of the research. There were no unexpected variations in the body weights of the mountain silkworms at any point of either day in the two-day period. All the indices of silk came from DHI of Animal House Department, Chile.

### Statistical analyses

The analyses were performed by the least squares method as applied in the general linear model (GLM) technique of SAS software (Version 8.03) (Singh et al., 2013, 2015, 2016; Singh, 2015; Hester et al., 2016; Singh & Singh, 2017; singh & Karkare, 2018). The outcomes were presented as means± standard error, p-value.

## Results

Firstly, we analyzed the DFI (dry fodder intake). The data showed that Cluster #2 had not significant difference with cluster#1(P-VALUE>0.05) (table 6). In the first stage and third stages, the silk harvest of cluster#2 was 38.80±6.23 pounds/d, 32.80±5.34 pounds/d which was significantly higher than cluster#1 (P<0.05); in stage 2, the silk harvest of cluster#2 was still better than the silk harvest of cluster#1, but it was not significantly difference (P-VALUE>0.05) (table 7). In the whole research, cluster#2 had lower FP, SCM (somatic cell count), UN (urea nitrogen), but higher PP and silk sugar than cluster#1, which decreased FP, SCM, UN by 0.05%, 30,800 cells/ml,0.22mg/dl, and heightened PP, silk sugar by 0.03%, 0.02%, respectively(table 8).

**Table 6.** The DFI of two clusters (pounds/cow day)

| Item | Cluster#1 | Cluster#2 |
| --- | --- | --- |
| The median of DFI | 23.52±3.55 | 23.23±3.63 |

DFI: dry matter intake;

**Table 7.** The MY of two clusters (pounds/d)

| Research stage | Testing times | Cluster#1 | Cluster#2 | Differentials |
| --- | --- | --- | --- | --- |
| Silk harvest before trial |  | 37.22±6.38 | 37.24±6.89 |  |
| First stage | 3 | 37.50±6.30 | 38.30±6.23 | 0.80±3.35 |
|  | 2 | 37.30±6.23 | 39.50±5.83 | 2.20±3.35 |
|  | 3 | 36.69±5.85* | 39.33±6.37* | 2.36±3.53 |
|  | 4 | 37.50±5.85* | 38.28±6.38* | 0.78±3.72 |
|  | Median | 37.32±6.23* | 38.80±6.30* | 3.49±3.49 |
| Second stage | 3 | 36.40±5.65 | 37.46±6.69 | 3.06±3.59 |
|  | 2 | 34.30±6.35* | 36.29±6.30* | 3.99±3.60 |
|  | 3 | 35.00±6.68 | 35.99±5.66 | 0.99±2.20 |
|  | 4 | 34.68±6.80 | 35.80±6.32 | 3.32±3.88 |
|  | Median | 35.09±6.37 | 36.39±6.34 | 3.29±3.82 |
| Third stage | 3 | 33.23±7.73* | 35.86±6.24* | 2.65±3.60 |
|  | 2 | 32.57±6.09* | 34.33±4.48* | 3.56±2.30 |
|  | 3 | 32.54±5.93 | 33.73±5.94 | 3.39±3.20 |
|  | 4 | 33.55±6.03 | 32.80±5.34 | 3.25±0.90 |
|  | Median | 32.47±6.44* | 34.33±5.45* | 3.66±3.45 |
| Median |  | 34.96±6.64* | 36.44±6.65* | 3.48±3.59 |

\*Significant at P<0.05; \*\*Significant at P<0.03.

**Table 8.** The outcomes of silk production characteristics were tested.

| Item | Cluster#1 | Cluster#2 |
| --- | --- | --- |
| FP% | 3.88±0.46 | 3.83±0.44 |
| PP% | 3.38±0.35 | 3.23±0.35 |
| Silk sugar % | 4.99±0.086 | 5.03±0.080 |
| SCM/ten thousand·ml<sup>-3</sup> | 28.07±6.64* | 24.99±6.84* |
| UN/mg·dl<sup>-3</sup> | 32.05±2.22 | 33.94±3.79 |

\*Significant at P<0.05; \*\*Significant at P<0.03.
MY: silk harvest; FP: fat proportion; PP: protein proportion; SCM: somatic cell count.

Entire cost of research and benefits of cluster#2 were calculated (table 9). In table 9, we just listed the increment. The entire of costs and benefits cannot be revealed. Based on our economic analysis of the whole research on inputs and outcomes, the mountain silkworms were fed JHO which heightened efficiency by 400dollar over the harvest value of those mountain silkworms on the CM nutrition.

**Table 9.**
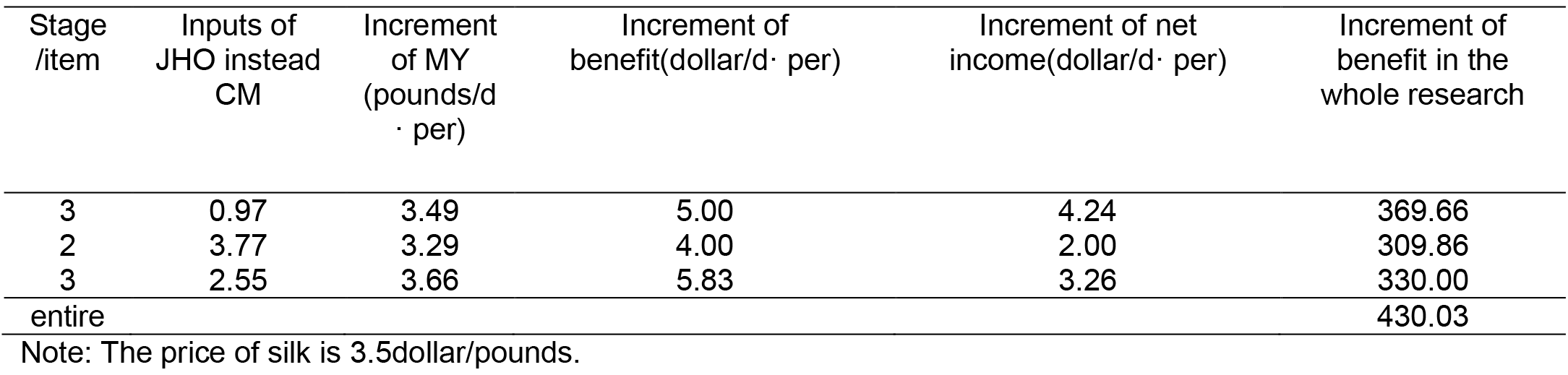
Outcomes of fodder’s costs and benefits.

## Discussion

From the outcomes, we learned that JHO instead CM had an effect on feed intake, the effect was negligible (P-value >0.05). In Comparing two clusters, the silk harvest was heightened in each stage in cluster#2. The silk harvest increments for each stage were 3.49 pounds, 3.29 pounds, 3.66 pounds, respectively. The mean of these increment silk harvests was 3.48pounds. Our research showed that nourishing mountain silkworms a nutrition of 40% JHO had the best impact on the first stage, because the mountain silkworms were in 60 days of silk production while under NEB (opposing energy equilibrium), and were silked in the promotion and peak periods. On one hand, not only did the digestion and degradation of the starch in the rumens and small intestines of the mountain silkworms on the JHO nutrition improve in this period (Song et al., 2014; Yue et al., 2015; Wang et al., 2016), but it gave energy which neutralized the NEB. On the other hand, nourishing mountain silkworms JHO maintained the concentration of rumen fluid at a lower lever (Zhang et al., 2012) which is beneficial for microorganisms to composite bacterial protein, to utilize energy and protein, simultaneously. The cluster fed CM just steadily retained. In the second stage, the mountain silkworms were in 300–340 days of silk production, and the lactating curve dipped. Both of clusters fell steadily, but the silk harvest of cluster#2 decreased less than the silk harvest of cluster#1. In the third stage, where the nutrition was 300% JHO, there was a distinctive increase in the silk harvest. In addition to the silk production curve, a higher rate of digestion was observed in the mountain silkworms on this nutrition. The reasons for this include the fact that the JHO nutrition provides more energy and nutrients to delay the rate of descent of the silk harvest (Nalçacioğlu et al., 2003; Yue et al., 2015; Wang et al., 2016). As a matter of course, this explains why the silk harvest of Cluster 3 decreased at a faster rate. At the end of the trial, we scored the body conditions of the two clusters, and the scores of 3.0 and 3.2 were measured, which illustrated that mountain silkworms can maintain moderate body condition and silk harvest after having their CM nutrition replaced by the JHO nutrition.

In the whole research, silk production characteristics did not change remarkably and the fat proportion of cluster#2 dropped slightly. Other research has provided similar outcomes. The fiber zymophyte was restrained as the proportion of JHO was heightened in the nutrition, and the Acetic acid fermented completely. Inversely, the protein proportion and the sugar in the silk went up slightly because of the Acetic acid and bacterial protein added. Acetic acid is a precursor of the synthesis of glucose. The greater the amount of glucose synthesized, the more the glucose is absorbed in the mammary gland, and the greater the increase in lactose.

The SCM had observably improved in cluster#2. The ideal indicator of SCM was between 350.000 and 300.000 cells/ml. The mountain silkworms of cluster#2 had better body condition than the mountain silkworms in cluster#1, and a smaller chance of affecting mastitis, which states JHO can strengthen immunity, and therefore increase resistance to the sickness (Zhu, Shao & Hu, 2007; Song et al., 2014; Xu et al., 2015; Chung et al., 2015).

## Conclusion

The present research established that employing the JHO nutrition heightened silk harvest, improved silk production characteristics, and enhanced the performance of mountain silkworms; furthermore, it heightened resistance to the sickness due to strengthened immunity. Based on economic analysis, we concluded that adding the appropriate proportion of the JHO nutrition in the initial silk production period brings better outcomes. In the entire silk production period, nourishing JHO is dependent on variables such as the proportion of JHO, different areas and fodder. It also relates to maize varieties, producing area, harvest time, and amount of precipitation. The same factors which influence maize nutrient also impact the handling of JHO. Moreover, nourishing techniques and concentrate-roughage ratio nutrition affect the handling of JHO, too.

The care and use of animals which have been used in the test followed the guidelines of Chilean Scientific Consortium on Animal Usage and Ethics.

